# Effect of charge on protein preferred orientation at the air–water interface in cryo-electron microscopy

**DOI:** 10.1101/2021.05.14.444152

**Authors:** Bufan Li, Dongjie Zhu, Huigang Shi, Xinzheng Zhang

## Abstract

The air–water interface (AWI) tends to absorb proteins and frequently causes preferred orientation problems in cryo-electron microscopy (cryo-EM). Here, we examined cryo-EM data from protein samples frozen with different detergents and found that both anionic and cationic detergents promoted binding of proteins to the AWI. By contrast, nonionic and zwitterionic detergents tended to prevent proteins from attaching to the AWI. This ability was positively associated with the critical micelle concentration of the detergent. The protein orientation distributions with different anionic detergents were similar and resembled that obtained without detergent. By contrast, cationic detergents gave distinct orientation distributions. The AWI is negatively charged and the absorption of cationic detergents to the AWI alters its charge. Our results indicates that proteins absorb to charged interface and the negative charge of the AWI plays an important role in absorbing proteins in the conventional cryo-EM sample preparation. According to these findings, a new method was developed to modify the charge distribution of the AWI by adding a very low concentration of anionic detergent. Using this method, the protein particles exhibited a more evenly distributed orientations and still absorbed to the AWI enabling them embedding in a thin layer of ice, which will benefit the cryo-EM structural determination.

## Introduction

After several decades of development, cryo-electron microscopy (cryo-EM) has become a powerful tool in structural biology. Recent rapid development of cryo-EM hardware and software has improved the resolution and expanded its application range in terms of the sizes of protein complexes. For cryo-EM, proteins need to be embedded in a thin layer of vitreous ice with an even distribution and random orientation. However, proteins often embed in vitreous ice with a preferred orientation, are only present in thick ice, or are sometimes denatured. These issues occur during cryo-EM sample preparation and are major barriers to high-resolution protein structure determination. It has been suggested that most of these phenomena are associated with emergence of an air–water interface (AWI) during blotting and absorption of proteins at the AWI before freezing.

Different methods have been proposed to separate proteins from the AWI. Detergents can occupy the AWI as the surface facing air is hydrophobic. Different detergents have been tested and work for some proteins(Chen et al., 2019). However, the screening of samples with different detergents is time-consuming and does not guarantee success. As an alternative method, grids have been coated with an ultra-thin carbon layer(Grassucci et al., 2007) (thickness: 3–5 nm) or graphene(Han et al., 2020; D’Imprima et al., 2019; Fan et al., 2019) to absorb proteins before they encounter the AWI. Ultra-thin carbon layers have a relatively strong background signal, which decreases the signal-to-noise ratio in the image and this is problematic when solving the structures of proteins that are 200 kDa or smaller. Graphene produces much less noise in cryo-EM images than ultra-thin carbon. However, the absorption of proteins on graphene is complicated and protein dependent. In addition, the interface between the film and water may cause different problems(Plevka et al., 2012). Collection of tilted images has also been used to compensate for preferred orientation(Tan et al., 2017). However, tilting reduces contrast because it increases the thickness of the ice and introduces additional beam induced motion.

During cryo-EM sample preparation, proteins are thought to move to the AWI through Brownian motion. If no other surface forces are present, theoretical calculations predict the duration of Brownian motion for a 100 kDa protein to reach the AWI will be 0.1 ms or less(Naydenova and Russo, 2017; Brune and Kim, 1993; Dubochet et al., 1988; Young et al., 1980). In this time, a protein could contact the AWI in a random orientation more than 1000 times before freezing, which gives ample opportunity for adsorption in a preferred orientation(Noble et al., 2018b; Sun, 2018; Naydenova and Russo, 2017). Thus, different spraying–freezing systems that allow for fast freezing after the formation of a thin film of water have been evaluated to freeze proteins before they are captured by the AWI in a preferred orientation(Kontziampasis et al., 2019; Ravelli et al., 2019; Rubinstein et al., 2019; Noble et al., 2018b). Some results have shown that the orientation distribution changes and becomes more even when the grid is frozen within 170 ms after formation of the thin film(Noble et al., 2018b). However, different systems have given contrasting results(Klebl et al., 2020) and more research is needed.

The mechanism behind absorption of proteins to the AWI needs to be elucidated and new methods need to be investigated to prevent absorption or reduce the preferred orientation problem. Here, we used tomography to study the distributions of proteins embedded in ice with different detergents added during freezing. Our data indicate that the main reason behind absorption of proteins at the AWI is its negative charge. According to this finding, we propose a new method to reduce the preferred orientation problem.

## Results and discussion

### The role of the critical micelle concentrations of nonionic and zwitterionic detergents in separating proteins from the AWI

The AWI tends to absorb most soluble proteins and causes preferred orientation problems(D’Imprima et al., 2019; Fan et al., 2019; Noble et al., 2018a; Noble et al., 2018b). By comparison, there are fewer orientation problems with membrane proteins. It has been suggested that detergents can occupy the AWI so that proteins are not in directly contact with it(Chen et al., 2019; Taylor and Glaeser, 2008; Quinn and Dawson, 1970). We studied the distributions of different membrane proteins in ice layers by cryo-tomography. With the detergents *n*-dodecyl-β-D-maltopyranoside (DDM), digitonin, *n*-dodecyl-*N,N*-dimethyl-amine-N-oxide (LDAO), and 3-[(3-cholamidopropyl)-dimethyl-ammonio]-1-propanesulfonate (CHAPS) (group 1) at a concentration above critical micelle concentration (CMC), the membrane proteins tended to stay between the two AWIs (Fig. 1). By contrast, except for in one case, group 2 detergents such as glycol diosgenin (GDN) and lauryl maltose neopentyl glycol (LMNG) resulted in the membrane proteins tending to form a single layer or two layers, indicating they absorbed to the AWI. Increasing the concentrations of these detergents did not affect the distribution of the proteins. It is worth noting that the molar ratios of the group 1 detergents calculated from the CMCs were higher than those in group 2 (Table S1). It is possible that the fraction of detergent above the CMC tends to form micelles and only the fraction of free detergent molecules below the CMC can bind to the AWI. Thus, a large CMC leads to more free detergent molecules that can bind to the AWI.

**Figure 1.**
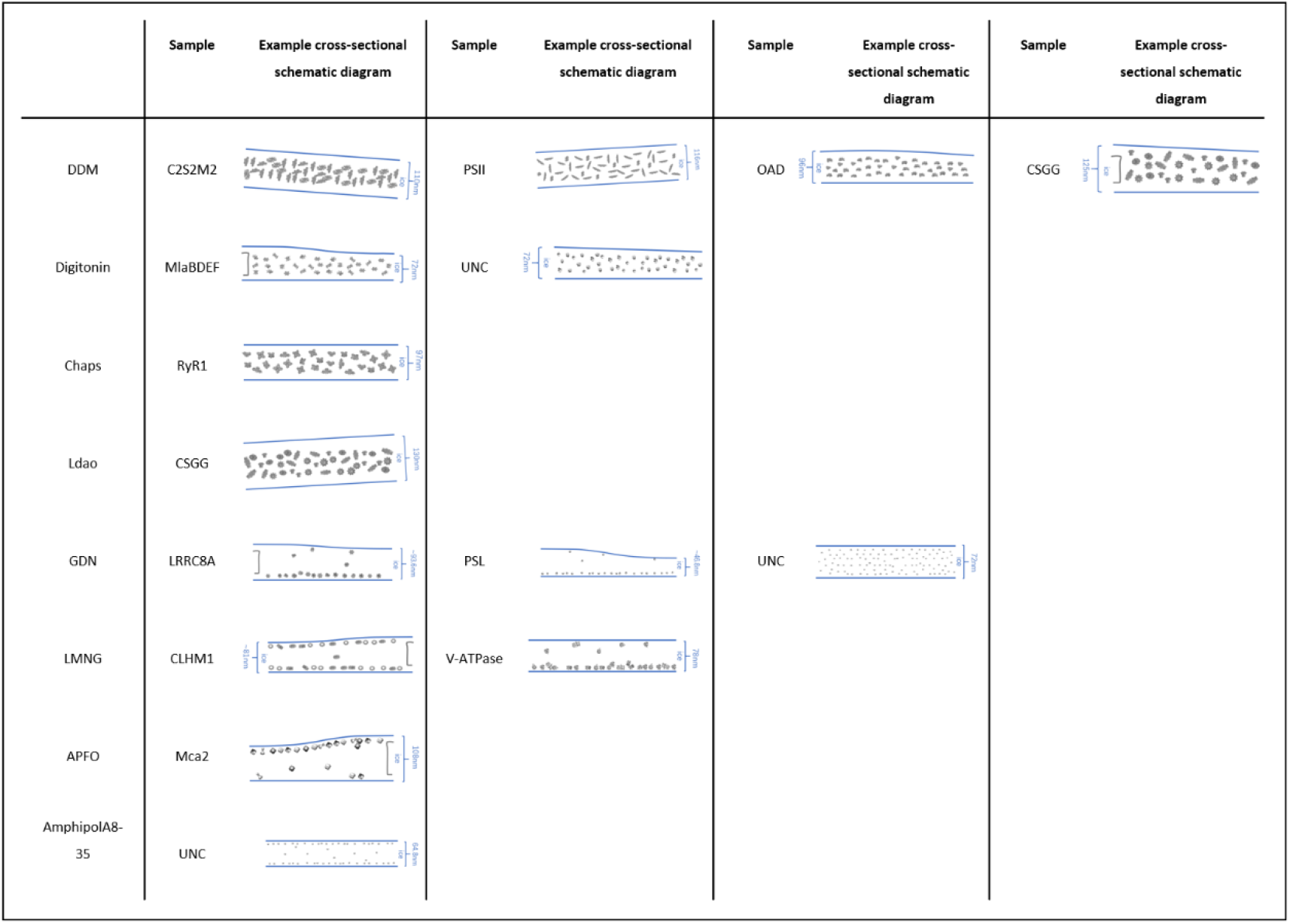
Cryo-tomography method for determining membrane protein behavior. The cross-sectional schematic diagrams of different membrane proteins embedded in vitreous ice were developed from the tomograms. The ice thicknesses are depicted accurately. Gray lines on schematic edges are the carbon film.

To achieve a reasonable concentration in the cryo-EM images, the concentration of proteins used for freezing in group 1 was much higher than that in group 2 (Table S2). These results are consistent with the phenomenon that the absorption of proteins to the AWI helps to enrich proteins(Klebl et al., 2020; Vinothkumar and Henderson, 2016; Rubinstein, 2007). We searched the Electron Microscopy Data Bank for the concentrations of different membrane proteins used in cryo-EM sample preparation. In most cases, the membrane protein concentrations in detergents with low CMCs were much lower than those of membrane proteins in detergents with high CMCs (Fig. S2 and Table S3), which agreed with our finding that the membrane proteins in detergents with low CMCs tended to absorb to and be enriched by the AWI.

For detergents in group 2, when the AWI is maintained for longer before freezing, it is still possible that the detergents may leave the micelles slowly and bind to the AWI. To investigate this, before freezing, we incubated two membrane proteins in 0.11% GDN and two membrane proteins in 0.11% LMNG for 5 min on grids. Interestingly, we found that the membrane proteins in GDN stayed between the two AWIs (Fig. S1), which indicated that GDN from the micelles may slowly diffuse to the AWI.

We also tested soluble proteins such as aldolase, β-galactosidase, and GDH to see whether detergents in groups 1 and 2 could keep soluble proteins away from the AWI. Our tomography results showed that without detergent these proteins stayed at a single AWI as a single layer or in two layers at both AWIs in the cryo-EM samples. These results agree with Noble’s observations(Noble et al., 2018a). Then, different detergents in both groups were used to occupy the AWI. When the group 2 detergents were used, soluble proteins stayed at the AWI, whereas they stayed between the two AWIs when the group 1 detergents were used (Fig. S3). Increasing the concentration of the group 2 detergent did not help the proteins leave the AWI. We also collected some single particle data for different detergents in different protein samples and our calculations showed that the orientations of soluble proteins that stayed between the two AWIs were random (Figs. S4 and S5).

### Orientation distributions for proteins absorbed to the AWI with cationic and anionic detergents

Ionic detergents were not considered in our above studies. These detergents have very high CMCs but are rarely considered as good solubilizers because they usually interfere with ion exchange separation(Healthcare and Healthcare, 2007; le Maire et al., 2000). To explore whether ionic detergents could prevent soluble proteins from absorbing to the AWI, 0.02% cetyltrimethylammonium bromide (CTAB) or cholesteryl hemisuccinate tris salt (CHS) was added to the buffer of aldolase, GDH, and 70S ribosome right before freezing. Interestingly, the tomographic data showed that all the proteins absorbed to the AWI in these experiments (Fig. S7).

To study the orientation of the protein absorbed to the AWI, we collected cryo-EM data for these samples. During blotting, the excess protein sample was blotted away from one AWI. We carefully loaded the cryo-EM grids into the microscope with this AWI facing up for data collection. During data processing, three-dimensional (3D) classification sometimes assigns redundant particles with preferred orientation to the wrong classes, which leads to a more uniform distribution of particle orientation in the right class. To preserve the original particle orientation distribution, 3D classification was excluded from our data processing. In addition, when there is a severe preferred orientation problem with particles in a cryo-EM data set, the densities of the reconstruction are distorted, which will affect determination of the particle orientations. Thus, we combined these particles with a good dataset collected in other experiments to ensure accurate determination of the particle orientations. We used a custom script to extract the orientation information of these particles (Fig. 2). For each protein, the orientation distribution of the sample with CTAB was quite different from that of the sample with CHS. Class-averaged images (Fig. S6) and orientation distribution maps (Fig. 2) showed that top views were dominant for aldolase and GDH with CHS, whereas side views were dominant when CTAB was added. With CTAB, 70S ribosome particles tended to stay in ice, thus a low concentration of 70S ribosome (80 nM) was used, which was much lower than the concentration used for freezing with CHS or without detergent. The dominant views of 70S ribosome in CHS differed from that in CTAB.

**Figure 2.**
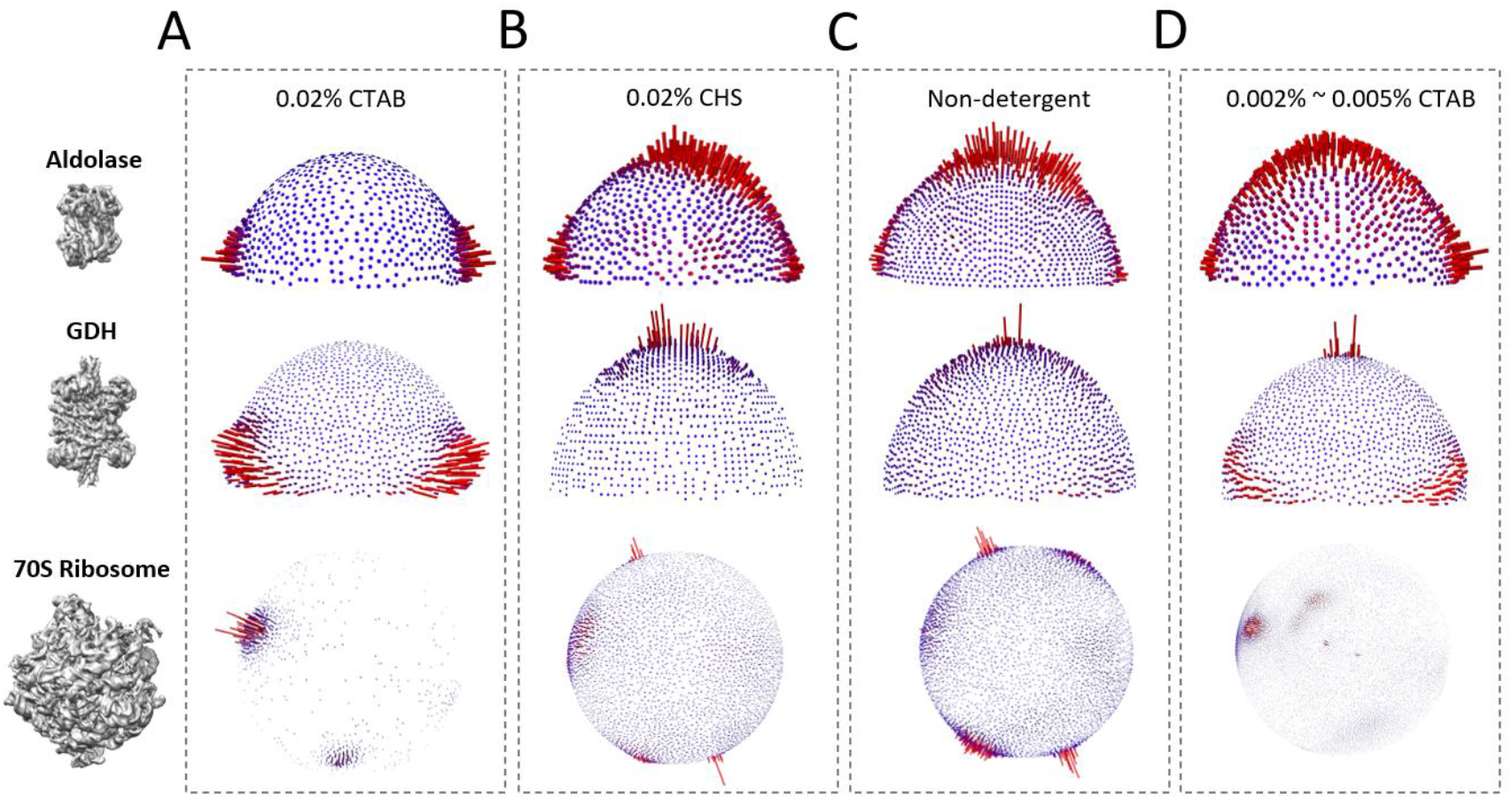
Orientation distributions obtained under different conditions. **A–D:** Orientation distribution spheres of the particles that contributed to the reconstructions on the left. The heights of the surface bars indicate the relative number of particles in a given orientation. **A:** Angular orientations of proteins in buffer with 0.02% CTAB. **B:** Angular orientations of proteins in buffer with 0.02% CHS. **C:** Angular orientations of proteins in conventional buffer without detergent. **D:** Angular orientations of proteins in buffer with 0.002%–0.005% CTAB.

Different types of cationic and anionic detergents, including dodecyl trimethyl ammonium chloride (DTAC) and sodium lauroyl sarcosine (SLS), were added to GDH to investigate the orientation distribution. Our results (Fig. S8) showed that the orientation distribution of the sample in DTAC was similar to that of the sample in CTAB, and the orientation distribution of the sample in SLS was similar to that of the sample in CHS. These results indicate that the positively or negatively charged hydrophilic head group determines the orientation distribution of the proteins.

### Negatively charged AWI absorbing proteins without detergent

Orientation distributions of different proteins in either cationic or anionic detergents were compared with those of the proteins without detergent. Interestingly, all proteins in anionic detergents exhibited orientation distributions similar to those of the same proteins without detergent (Fig. 2B and C). If both anionic and cationic detergents occupied the AWI as nonionic and zwitterionic detergents do(Glaeser et al., 2016; Vos et al., 2008; Colloid and Science, 2004; le Maire et al., 2000), the AWI with anionic detergents would be negatively charged. Because the interfaces with anionic detergents exhibit orientation distributions similar to the AWI without detergent, this indicates that the AWI without detergent is also negatively charged. Indeed, Masayoshi and Drzymala et. al directly measured the surface potentials of microbubbles in aqueous solutions and found that the AWI was negatively charged(Chaplin, 2009; Takahashi et al., 2002; Drzymala et al., 1999).

For samples with cationic detergents, the AWI is positively charged, which leads to different orientation distributions. However, it is still possible that cationic detergents bind to the protein and affect the orientation. First, if ionic detergents bind to proteins, different types of ionic detergents may bind to different parts of the protein and cause different orientation distributions. This does not agree with our finding that the orientation distributions only depend on the type of charge on the hydrophilic head group. To further exclude this possibility, we performed buffer exchange on the cation detergent sample to remove free detergent molecules. In this case, the cationic detergent could not occupy the AWI in the cryo-EM sample, but it could still associate with proteins. The protein orientation distribution in this sample resembled that in the non-detergent sample (Fig. S9). Thus, it is very possible that the cationic detergent makes the AWI positively charged, which results in a change in protein orientation.

Our results indicate that it is the electric charge rather than the hydrophobic nature of the AWI that plays an important role in determining the orientation distribution.

### Modifying the charge property of AWI to reduce the preferred orientation

Cationic detergents changed the original negative charge of the AWI to a positive charge and altered the orientation of proteins absorbed to the interface. In this case, it was possible to reach an equilibrium point when a certain concentration of cationic detergent made the net charge of the interface close to zero. We tried to find this point by screening different concentrations of cationic detergents and finding a concentration that resulted in the proteins leaving the AWI. However, we were not able to find such a concentration, which was probably because the range for the equilibrium point was very narrow. However, with CTAB concentrations between 0.002% and 0.005%, we found a new orientation distribution representing a mixture of orientation distributions from samples with cationic detergents and without detergent (Figs. 2D and 3). Thus, our results show that adding a certain concentration of cationic detergent introduces new protein orientations, which may help to give a more uniform distribution.

**Figure 3.**
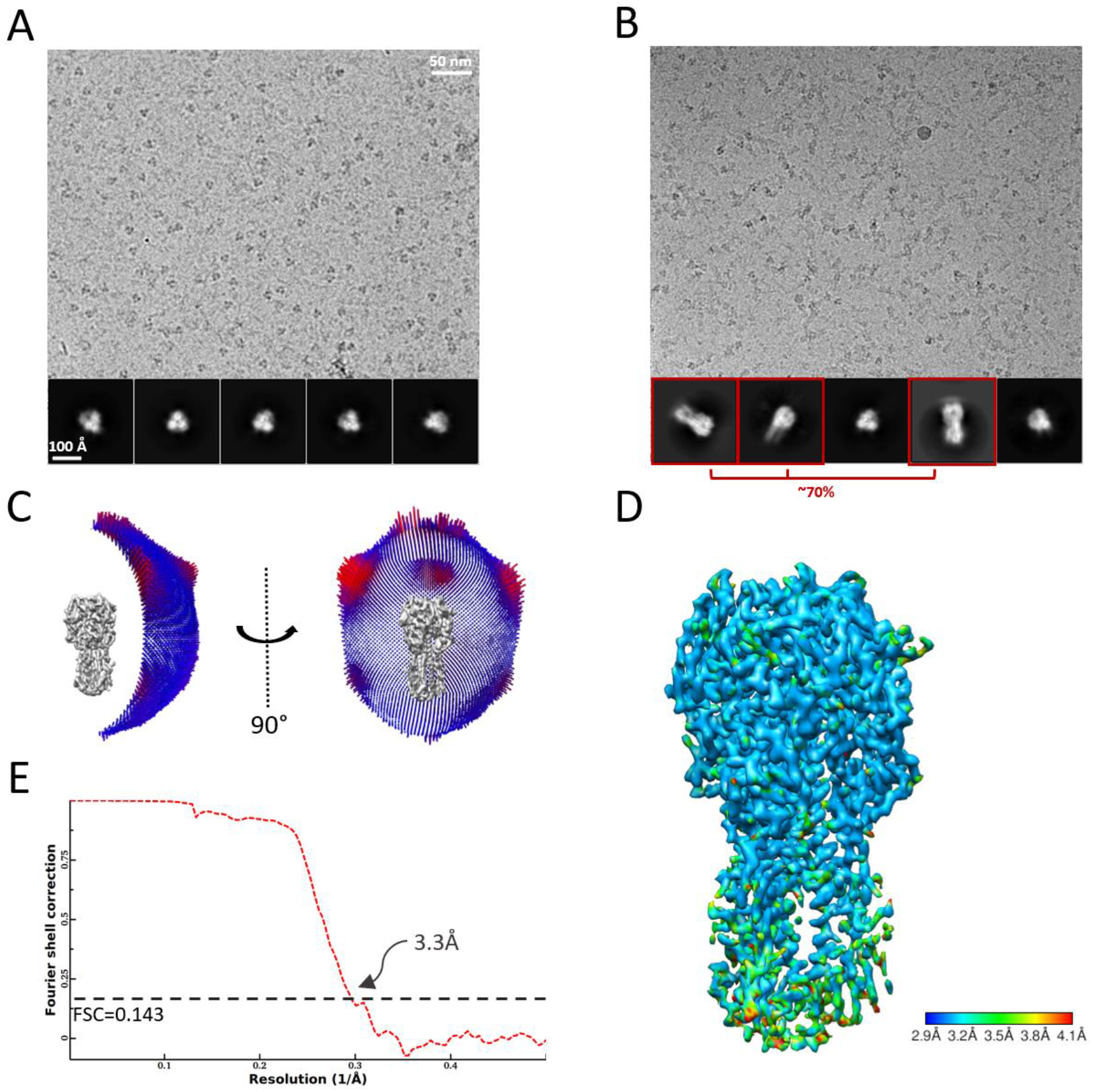
Single particle image processing flow-chart of the hemagglutinin dataset. **A:** Representative high-magnification image and 2D class averages of HA in conventional buffer without detergent. **B:** Representative high-magnification image and 2D class averages of HA in buffer with CTAB. B–E: The results from different image processing steps, including two-dimensional (2D) classification (B), final 3D auto-refining (C), the local resolution map (D), Euler angle distribution (C), directional FSC profile (E).

To quantitatively measure the improvement of orientation distribution, the orientation efficiency, *E*_od_ as defined by Naydenova et. al., were calculated[9] on different proteins (Table S6). For aldolase, GDH and influenza hemagglutinin (HA), the *E*_od_ with low concentration of CTAB is better than that from proteins frozen by the conventional freezing method. However, since using conventional freezing method, HA has severe preferred orientation problem, thus the value of *E*_od_ with low concentration of CTAB exhibits a large increase on HA sample. However, it has been shown that 70S ribosomes prefer to absorbed to positively charged interface, thus, in our dataset with low concentration of CTAB, ribosomes absorbed to surface with CTAB were dominant. If we randomly pick the same number of 70S ribosome particles from the dataset by conventional freezing and combine these particles with the dataset with low concentration of CTAB, the *E*_od_ is also improved as shown in Table S6.

Detergent addition has been proposed as an approach to prevent absorption of soluble proteins at the AWI(Chen et al., 2019; Glaeser and Han, 2017). Our approach is totally different because the proteins still absorb to the AWI and tend to be present in thin ice, so that the AWI enriches the protein concentration. The concentration of detergent used in our approach is lower than that required for the previous method, which is good for retaining contrast. For example, top views are dominant in regular sample preparation of HA trimer protein, which distorts densities in the reconstruction(Tan et al., 2017). By comparison, when we added CTAB at 0.005% to HA protein, the proportion of side views increased to 70% (Fig. 3A and B) and we obtained a 3.3 Å reconstruction without density distortion (Fig. 3C, D and E).

### Modeling of protein absorption by the AWI

The surface charges at the AWI are shielded by the salt solution, which forms a potential well between the surface and where the charges are fully shielded. The potential well can trap proteins and rotate them to a preferred orientation so that the local electrostatic potential of the protein or partial protein in the potential well is minimized. A model with proteins contacting the AWI through Brownian motion has been proposed(Klebl et al., 2020; Glaeser and science, 2018; Noble et al., 2018b; Sun, 2018; Glaeser and Han, 2017; Naydenova and Russo, 2017; Taylor and Glaeser, 2008). A simple calculation shows that it takes approximately 0.1 ms for a 100 kDa protein to reach the AWI(Naydenova and Russo, 2017; Brune and Kim, 1993; Dubochet et al., 1988; Young et al., 1980). If the orientation of the protein is random when the protein contacts the AWI, the protein will leave the AWI and rejoin the interface in another round of Brownian motion. The protein may undergo tens to thousands rounds of Brownian motion before it is captured by the AWI in the preferred orientation, which could take several milliseconds. However, in our model (Fig. 4), the potential well would help the protein rotate to the preferred orientation, and the AWI could capture the protein after a single round of Brownian motion, which would take only approximately 0.1 ms. This is consistent with the recent finding(Klebl et al., 2020) that rapid freezing of a protein within 6 ms still causes absorption of proteins to the AWI with preferred orientations.

**Figure 4.**
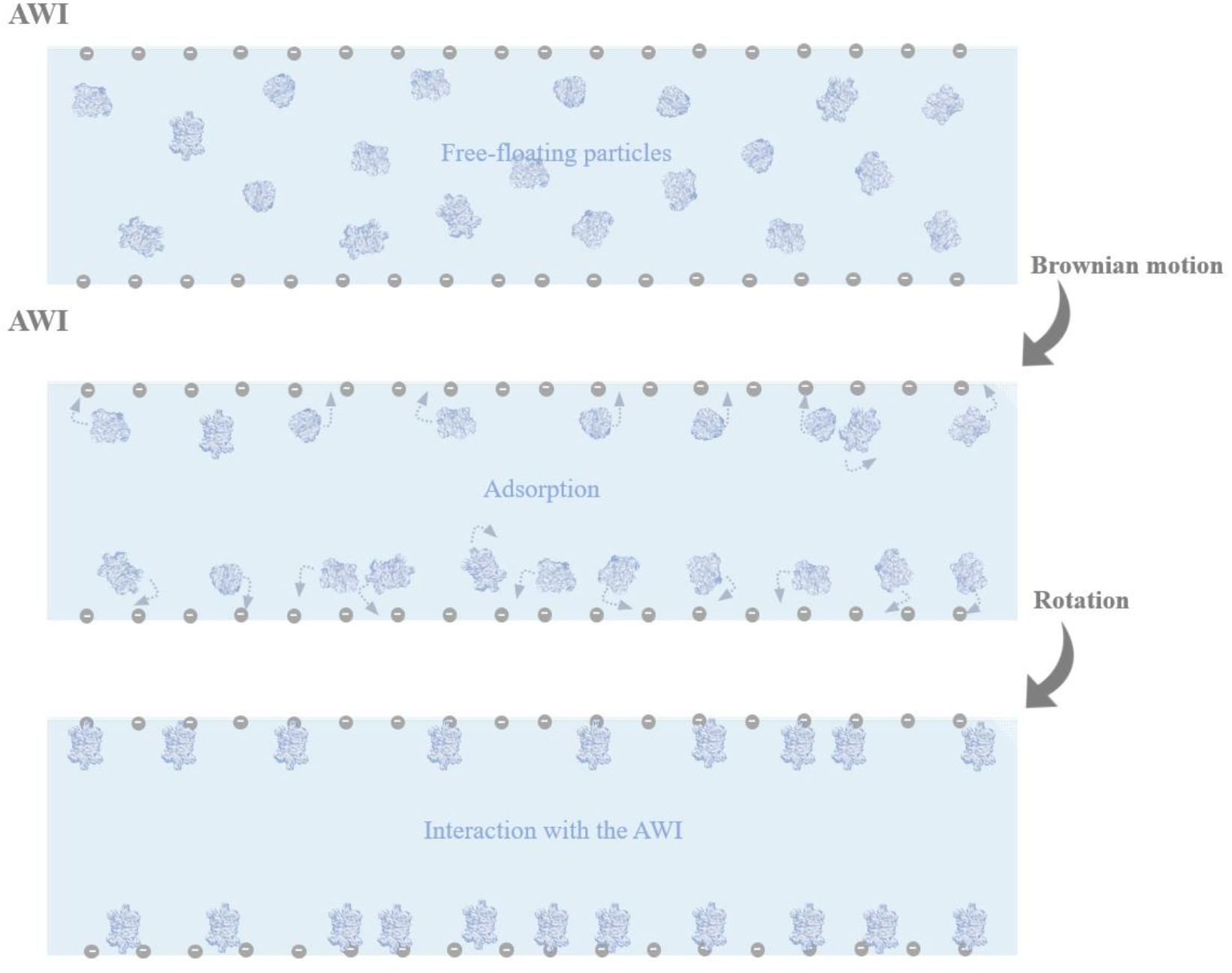
Proposed model of protein–AWI interactions.

## Conclusions

We found that nonionic and zwitterionic detergents with higher CMC values can prevent both membrane proteins and soluble proteins from binding the air-water interface, thereby reducing the preferred orientation problems. Contrary, ionic detergents tend to adsorb protein particles on the air-water interface. The orientation distribution of proteins in anionic detergents was quite different from cationic detergents, but similar to the buffer without detergent. These results revealed that the negative charge on AWI plays the major role in absorbing proteins. According to this finding, we added very low concentration of cationic detergent to protein buffer to modify the negatively charged AWI and we found better orientation distribution. Using this method, the proteins were present in thin ice and absorbed to the AWI with a more evenly distributed orientations, which retains the contrast of image.

## Materials and methods

### Sample preparation

For membrane proteins, most of the grids characterized were prepared using conventional techniques as determined by the sample owner. Generally, 3 μL of the sample was added to a plasma-cleaned (Solarus, Gatan) holey grid and plunge frozen in liquid ethane using a Vitrobot (Gatan) or EM GP (Leica).

Aldolase from rabbit muscle (Sigma–Aldrich) was solubilized in buffer (20 mM HEPES [pH 7.5], 50 mM NaCl) at approximately 3 mg/mL and then further purified by size-exclusion chromatography using a Superdex-200 column (GE Healthcare) equilibrated in solubilization buffer. Aldolase was concentrated immediately before cryo-EM grid preparation. The aldolase concentration used for cryo-EM sample preparation depended on the type of detergent. For samples without detergent, with nonionic and zwitterionic detergents of low CMCs, and with ionic detergents, the aldolase concentration was approximately 1.6 mg/mL, and that for samples with nonionic and zwitterionic detergents of high CMCs was approximately 6 mg/mL.

β-Galactosidase (catalog #G5635, Sigma–Aldrich) was subjected to gel-filtration chromatography on a Superdex-200 size-exclusion column connected to an ÄKTA FPLC apparatus (GE Healthcare). The elution buffer contained 25 mM Tris (pH 8), 50 mM NaCl, 2 mM MgCl_2_, and 1 mM TCEP. The β-galactosidase concentration used for cryo-EM sample preparation without detergent, with nonionic and zwitterionic detergents of low CMCs was approximately 2.0 mg/mL, and that with nonionic and zwitterionic detergents of high CMCs was approximately 10.0 mg/mL.

GDH (catalog #G2626, Sigma–Aldrich) was dialyzed in buffer (100 mM potassium phosphate, pH 6.8) overnight before purification by gel filtration. The GDH concentration used for cryo-EM sample preparation without detergent, with nonionic and zwitterionic detergents of low CMCs, and with ionic detergents was approximately 3.6 mg/mL. The GDH concentration with nonionic and zwitterionic detergents of high CMCs was approximately 12 mg/mL.

*Escherichia coli* 70S ribosome, a gift from Professor Qin’s laboratory, was diluted to a concentration of approximately 80 nM. The concentration of *E. coli* 70S ribosome in CTAB for cryo-EM sample preparation was approximately 80 nM, and that without detergent or in CHS was approximately 0.5 μM.

Hemagglutinin was prepared as described by Tan et al(Tan et al., 2017).

Sample aliquots of approximately 3 μL were applied to separate glow-discharged NiTi foil R1.2/1.3(Au) grids. After blotting for 3.5 s, the grids were flash plunged into liquid ethane at approximately 180 °C by a Vitrobot (Gatan).

### Single-particle cryo-EM data collection

Datasets of aldolase without detergent and aldolase in the detergents LMNG, CHAPS, DDM, CHS, and CTAB were collected using a Talos Arctica (FEI) equipped with a direct detector K2 summit (super resolution mode) and GIF quantum energy filter (energy width of 20 eV) at a pixel size of 1.0 Å. The corresponding dose rate was approximately 9.5 e^−^/Å^2^/s and the exposure time was 6.4 s. The images were fractionated into 32 frames. The defocus range of the collected micrographs was −1.5 μm to −2.0 μm.

Under the same conditions, we also collected datasets for the following: GDH both without detergent and in CHS, CTAB, DTAC, and SLS; and β-galactosidase both without detergent and in CHAPS, DDM, CHS, and CTAB.

Datasets of *E. coli* 70S ribosome both without detergent and in CHS and CTAB were collected using the Talos Arctica or a Titan Krios G2 (FEI) equipped with a direct detector K2 summit (super resolution mode) at dose rates of approximately 9.34 e^−^/Å^2^/s, 9.5 e^−^/Å^2^/s, and 9.89 e^−^/Å^2^/s. The exposure times were 11.2 s, 6.4 s, and 11.2 s, yielding binned pixel sizes of 1.32 Å, 1.0 Å, and 1.36 Å, respectively. The images were fractionated into 32 frames. The defocus range was −1.5 μm to −2.0 μm.

The HA trimer sample was imaged on the Talos Arctica equipped with a direct detector K2 summit (super resolution mode) and GIF quantum energy filter (energy width of 20 eV) at 1.0 Å/pixel in counting mode. The dose was approximately 60 e^−^/Å^2^ across 32 frames, which gave a dose rate of approximately 9.5 e^−^/pixel/s. The defocus range was −1.5 μm to −2 μm. A total of approximately 600 raw micrographs were collected.

### Processing of single-particle cryo-EM data

The frames were corrected by MotionCor2. The defocus parameters were estimated by CTFFIND4. Particles were auto-picked by and iteratively two-dimensionally classified by Relion 3.0, yielding featured two-dimensional (2D) averages. After discarding false particles in reference-free 2D and 3D classifications, the particles were kept for further refinement, and the resolution of the final 3D reconstruction was estimated using the FSC 0.143 criterion. The 3D renderings of the maps and models were created in UCSF-Chimera.

### Collection and processing of cryo-ET tilt-series data

Most tilt series were collected on Talos 200 microscopes (Thermo Fisher Scientific) with a Ceta camera using SerialEM at a dose rate of approximately 5 e^−^/Å^2^/s and a total accumulated dose of approximately 130 e^−^/Å^2^. The magnification for collection of most of the tilt-series data was 57 000, and the calibrated pixel size was 1.79 Å. Most tilt series were collected bi-directionally with a tilt range of −45° to 46° and a tilt increment of 3.5°. All tilt series were acquired at a defocus of about −8.0 μm. The tilt series were aligned and reconstructed with a binning factor of four in the IMOD package. Most tilt series were coarsely aligned, but some were manually aligned if necessary, and CTF correction was not performed.

Small ice crystals from atmospheric contamination on the surface of the vitrified layer were used to locate the two AWI, and the distance between the two interfaces was measured. The measuring error of the ice thickness was several nanometers.

## Acknowledgements

We thank Y. Rong, Y.L. Han, Dr. L. Xiao, C.P. Zhou, W.X. Yang, C.Y. Wang, X.P. Wang, L. Si, Dr. X.D. Su, B.H. Zou, Prof. K. Zhang, Prof. M. Li, Prof. Y.H. Huang and Prof. Z.F. Liu at the Institute of Biophysics, Chinese Academy Sciences, Beijing, China for preparing the membrane protein samples. We thank Dr. X.T. Cao at the Institute of Biophysics, Chinese Academy Sciences, Beijing, China for preparing E. coli 70S ribosome sample. We thank L.F. Kong at the Institute of Biophysics, Chinese Academy Sciences, Beijing, China for cryo-EM data storage and backup. Cryo-EM data collection was carried out at the Center for Biological Imaging, Core Facilities for Protein Science at the Institute of Biophysics (IBP), Chinese Academy of Sciences (CAS). We thank X.J. Huang, L.H. Chen, F. Sun, and other staff members at the Center for Biological Imaging (IBP, CAS). The project was funded by the National Key R&D Program of China (2017YFA0504700), the Natural Science Foundation of China (31930069) the Strategic Priority Research Program of the Chinese Academy of Sciences (XDB37040101) and the Key Research Program of Frontier Sciences at the Chinese Academy of Sciences (ZDBS-LY-SM003).

## Figures

**Table S1.** Properties of detergents used in in in this study. The physical properties including CMC and molecular weight are important when optimizing cryo-specimen conditions. Red box represents the group 1, and blue box represents the group 2.

| Detergent | CMC (mM) | CMC (%w/v) | Molecular Weight (Da) | Type |
| --- | --- | --- | --- | --- |
| **DDM**<br>n-Dodecyl-β-D-Maltopyranoside | ~ 0.17 | 0.0087% | 510.6 | Nonionic |
| **Digitonin** | <0.5 | 0.061% | 1229.31 | Nonionic |
| **LDAO**<br>n-Dodecyl-N, N-Dimethylamine-N-Oxide | ~ 1-2 | 0.023% | 229.4 | Zwitterionic |
| **CHAPS**<br>3-[(3-Cholamidopropyl)-Dimethyl-Ammonio]-1-Propanesulfonate | ~ 8 | 0.49% | 614.9 | Zwitterionic |
| **FC10**<br>Fos-Choline-10 | ~ 11 | 0.35% | 323.4 | Zwitterionic |
| **Cymal-3**<br>3-Cyclohexyl-1-Butyl-β-D-Maltoside | ~ 34.5 | 1.6% | 466.5 | Nonionic |
| **GDN**<br>Glycol Diosgenin | ~0.018 | 0.0021% | 1165.31 | Nonionic |
| **LMNG**<br>Lauryl Maltose Neopentyl Glycol | ~0.01 | 0.001% | 1005.19 | Nonionic |
| **CHS**<br>Cholesteryl Hemisuccinate Tris Salt | N/A | N/A | 607.9 | Anionic |
| **CTAB**<br>(1-Hexadecyl) Trimethyl Ammonium Bromide | ~0.92 | 0.0335% | 364.446 | Cationic |
| **SLS**<br>Sodium Lauroylsarcosine | 0.871 | 0.0256% | 293.39 | Anionic |
| **DTAC**<br>Dodecyl Trimethyl Ammonium Chloride | 20 | 0.53% | 263.89 | Cationic |

\*N/A = Not available

**Table S2.**
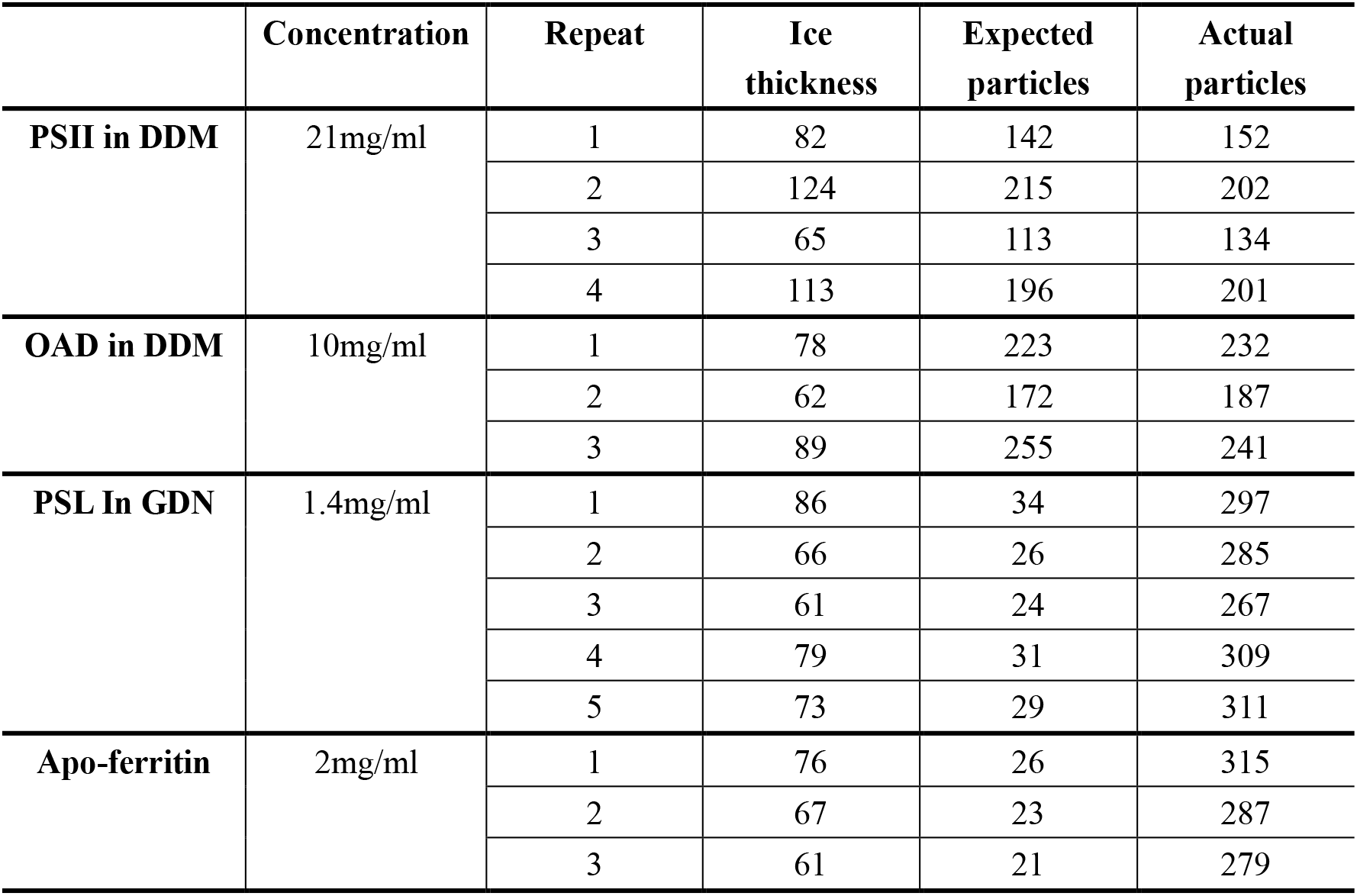
A summary of expected and actual particles distribution on one cryo-EM micrograph.

|  | Concentration | Repeat | Ice thickness | Expected particles | Actual particles |
| --- | --- | --- | --- | --- | --- |
| **PSII in DDM** | 21mg/ml | 1 | 82 | 142 | 152 |
|  |  | 2 | 124 | 215 | 202 |
|  |  | 3 | 65 | 113 | 134 |
|  |  | 4 | 113 | 196 | 201 |
| **OAD in DDM** | 10mg/ml | 1 | 78 | 223 | 232 |
|  |  | 2 | 62 | 172 | 187 |
|  |  | 3 | 89 | 255 | 241 |
| **PSL In GDN** | 1.4mg/ml | 1 | 86 | 34 | 297 |
|  |  | 2 | 66 | 26 | 285 |
|  |  | 3 | 61 | 24 | 267 |
|  |  | 4 | 79 | 31 | 309 |
|  |  | 5 | 73 | 29 | 311 |
| **Apo-ferritin** | 2mg/ml | 1 | 76 | 26 | 315 |
|  |  | 2 | 67 | 23 | 287 |
|  |  | 3 | 61 | 21 | 279 |

**Table S3.**
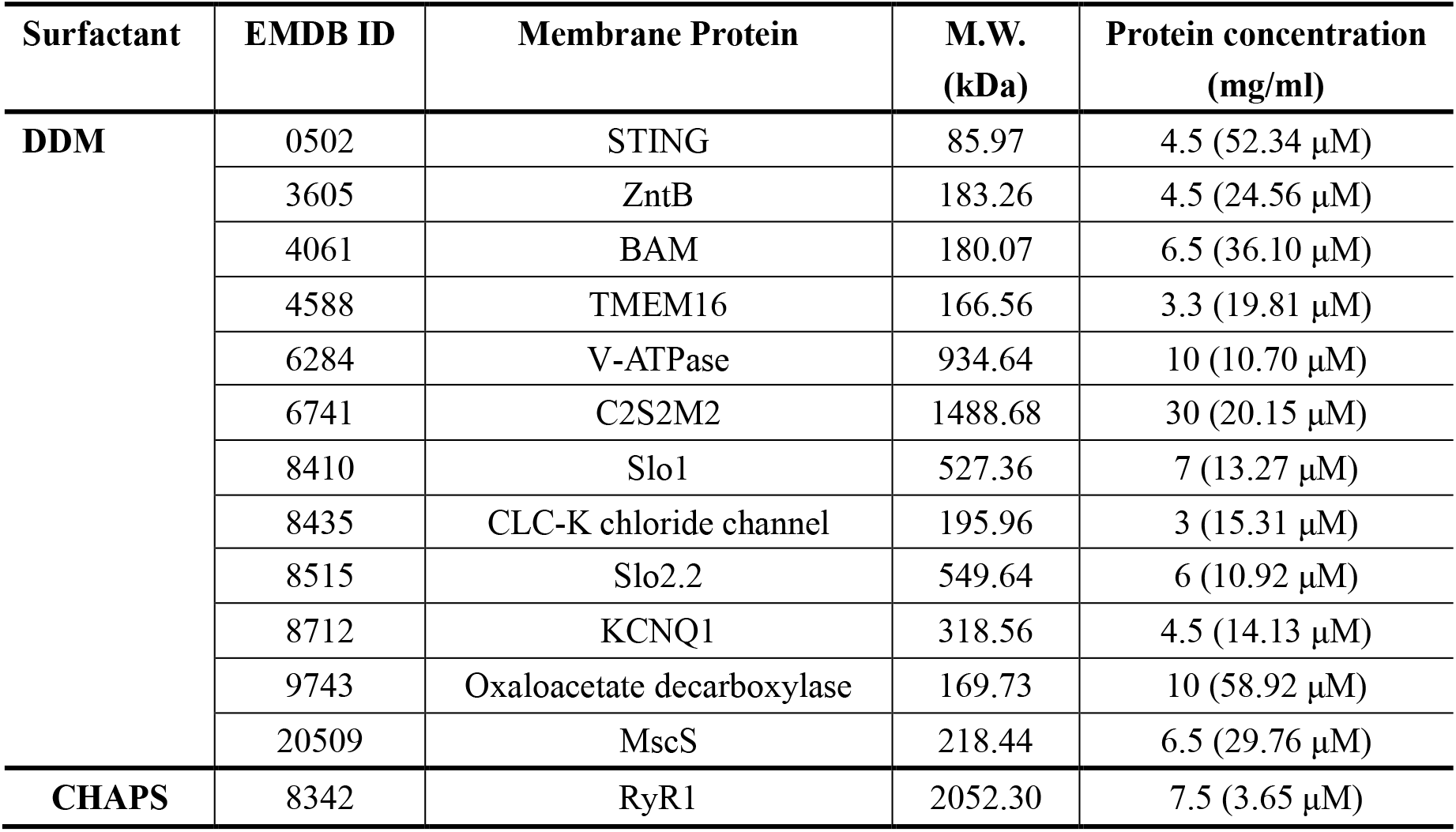
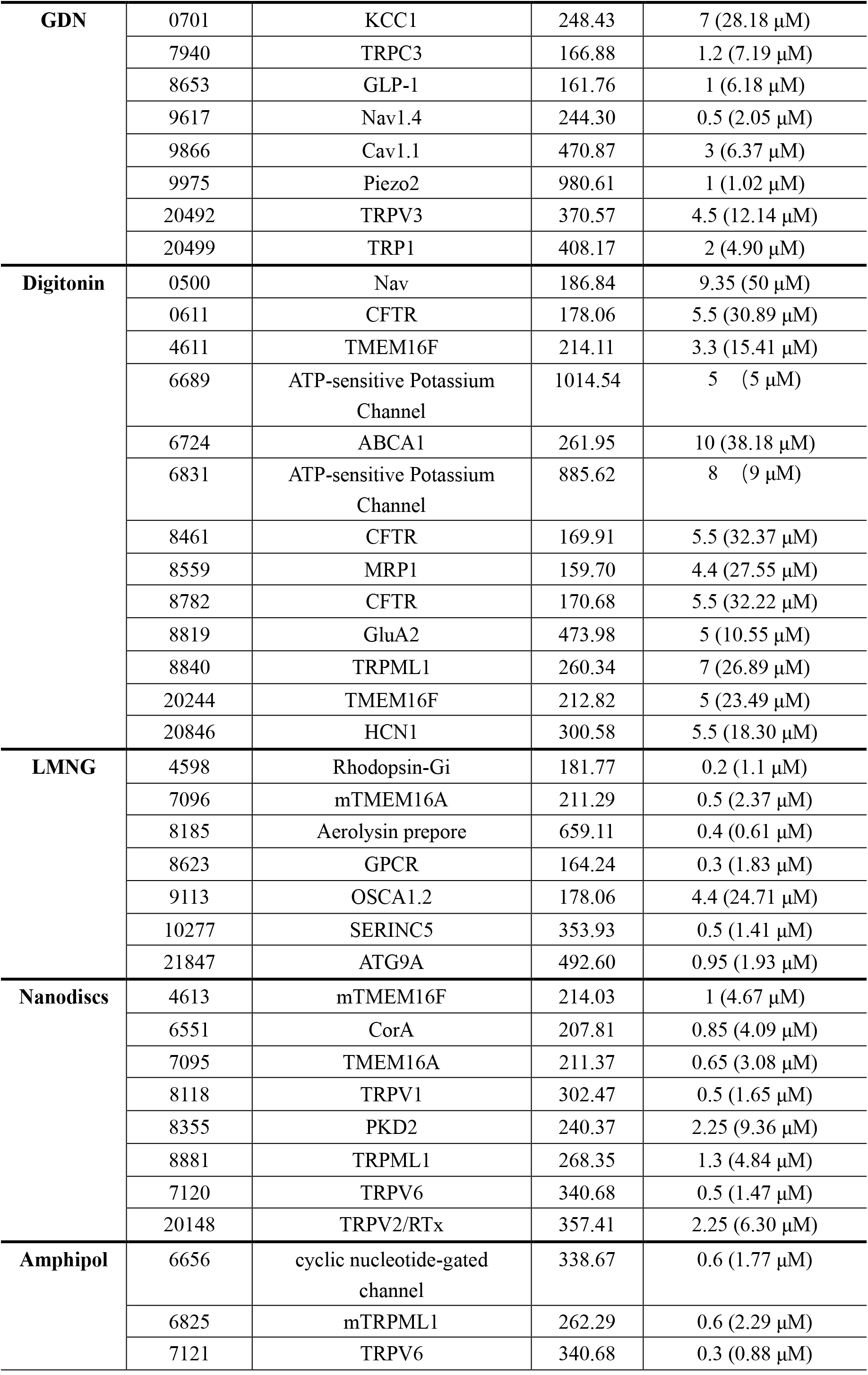
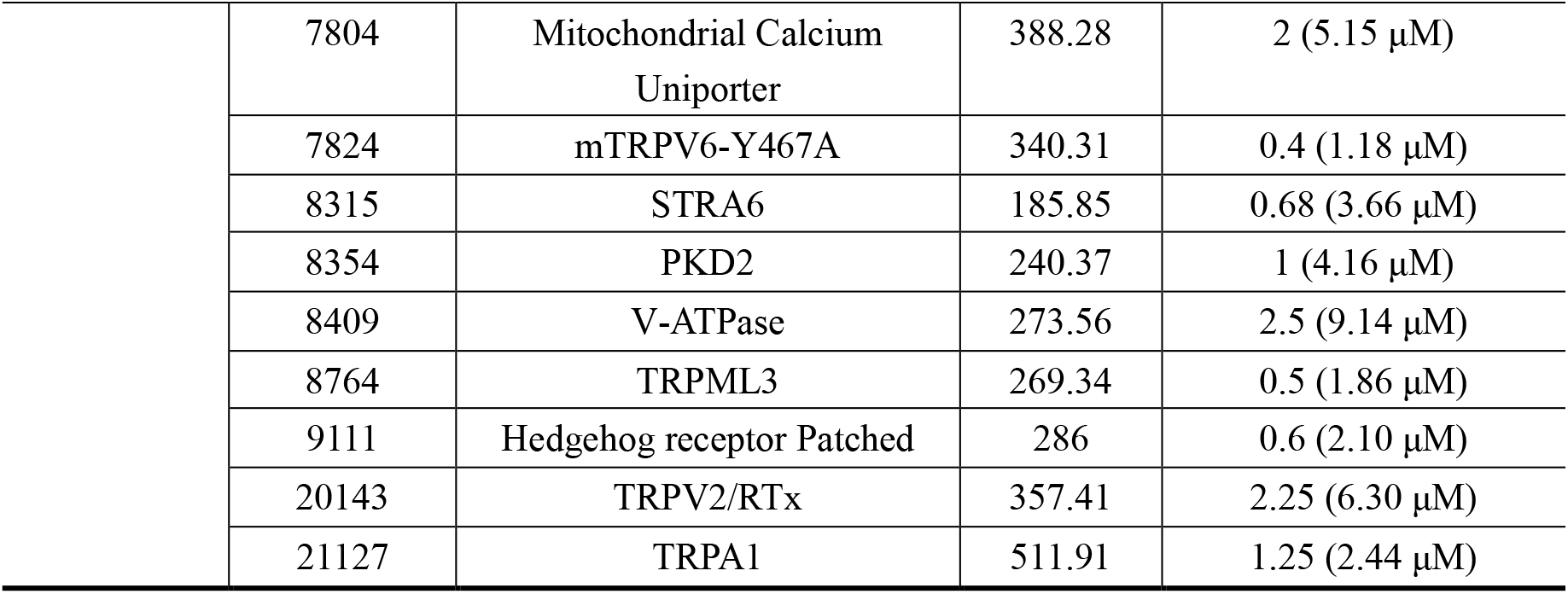
A summary of concentration and other attributes of membrane protein complexes determined by single-particle cryo-EM. Statistics are derived from the EMDB databank.

**Table S4.** Microscope parameters for collection of cryo-ET data.

|  |  |
| --- | --- |
| **Microscope** | Talos F200C |
| **Magnification** | 57,000 |
| **Camera** | Ceta |
| **Voltage (kV)** | 200 |
| **Electron dose per image ($\text{e}^-/\text{\AA}^2$)** | 5 |
| **Fractions per image** | 1 |
| **maximum/minimum tilt (increment)** | -45°, +46° (3.5°) |
| **Defocus range ($\mu\text{m}$)** | -8 |
| **Pixel size ($\text{\AA}$)** | 1.79 |

**Table S5.** Data collection parameters for SPA datasets.

|  |  |  |
| --- | --- | --- |
| **Microscope** | Talos Arctica | Titan Krios G2 |
| **Camera** | K2 | K2 |
| **Voltage (kV)** | 200 | 300 |
| **Energy width (eV)** | 20 | 20 |
| **Total electron dose ($\text{e}^-/\text{\AA}^2$)** | 60 | 60 |
| **Exposure time (s)** | 6.4 or 11.2 | 11.2 |
| **Number of frames** | 32 | 32 |
| **Defocus range ($\mu\text{m}$)** | -1.5 ~ -2 | -1.5 ~ -2 |
| **Pixel size ($\text{\AA}$)** | 1.0 or 1.32 | 1.36 |

**Table S6.** The orientation efficiency (*E*_od_) for aldolase, GDH, 70S Ribosomes and HA without detergent and with 0.002%∼0.005% CTAB.

|  | **Aldolase** | **GDH** | **70S**<br>**Ribosomes** | **HA** |
| --- | --- | --- | --- | --- |
| **$E_{od}$ of 0.002%~0.005%**<br>**CTAB** | 0.82 | 0.85 | 0.55 (0.74) | 0.87 |
| **$E_{od}$ of Non-detergent** | 0.77 | 0.71 | 0.69 | 0.16 |

**Figure S1.**
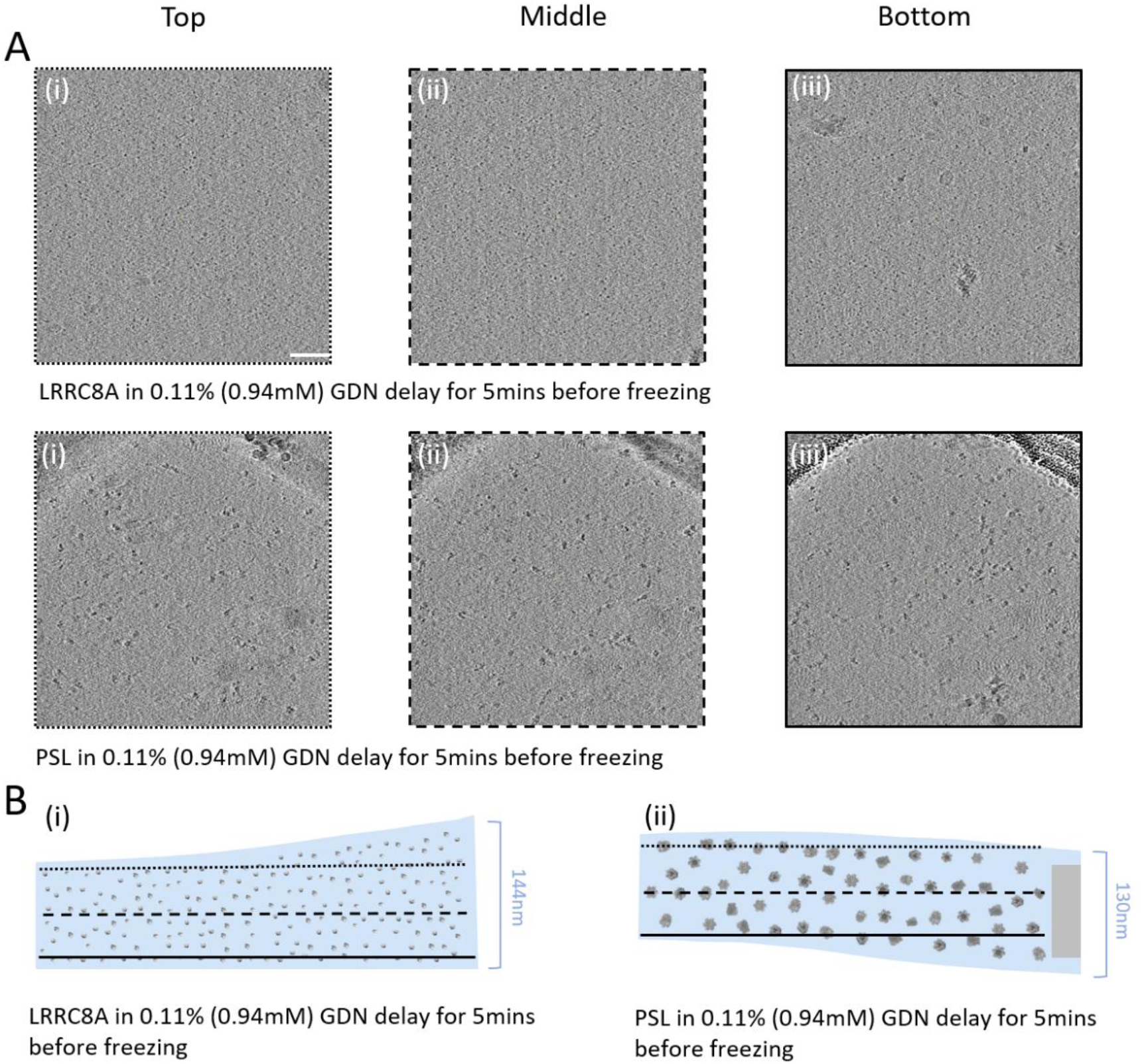
Incubated the membrane proteins in the detergents of GDN before freezing. A: Surface tomographic cross-section of the vitreous ice layer. (Left Panel) Top tomographic cross-section. (Middle Panel) Middle tomographic cross-section. (Right Panel) Bottom tomographic cross-section. B: Schematic diagram of particle distribution in ice. Thickness of ice is indicated on the right. Different lines represent relative depths through tomographic reconstructions.

**Figure S2.**
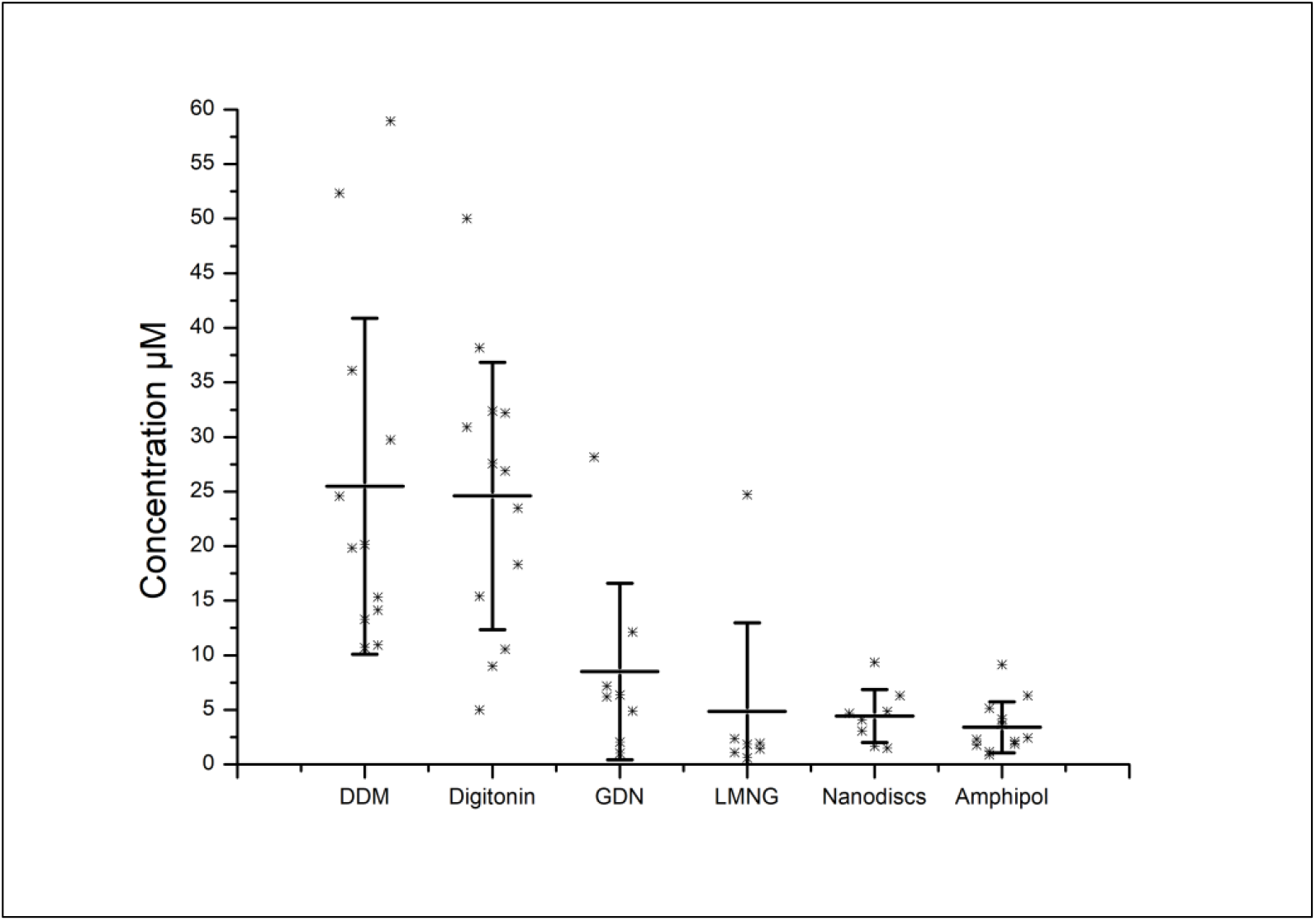
The concentrations of different membrane proteins used in cryo-EM sample preparations. The sample concentrations of membrane proteins in DDM, digitonin, GDN, LMNG, Nanodiscs and A835 were counted, statistics are derived from the EMDB databank. DDM and Digitonin requires a much higher sample concentration than the others.

**Figure S3.**
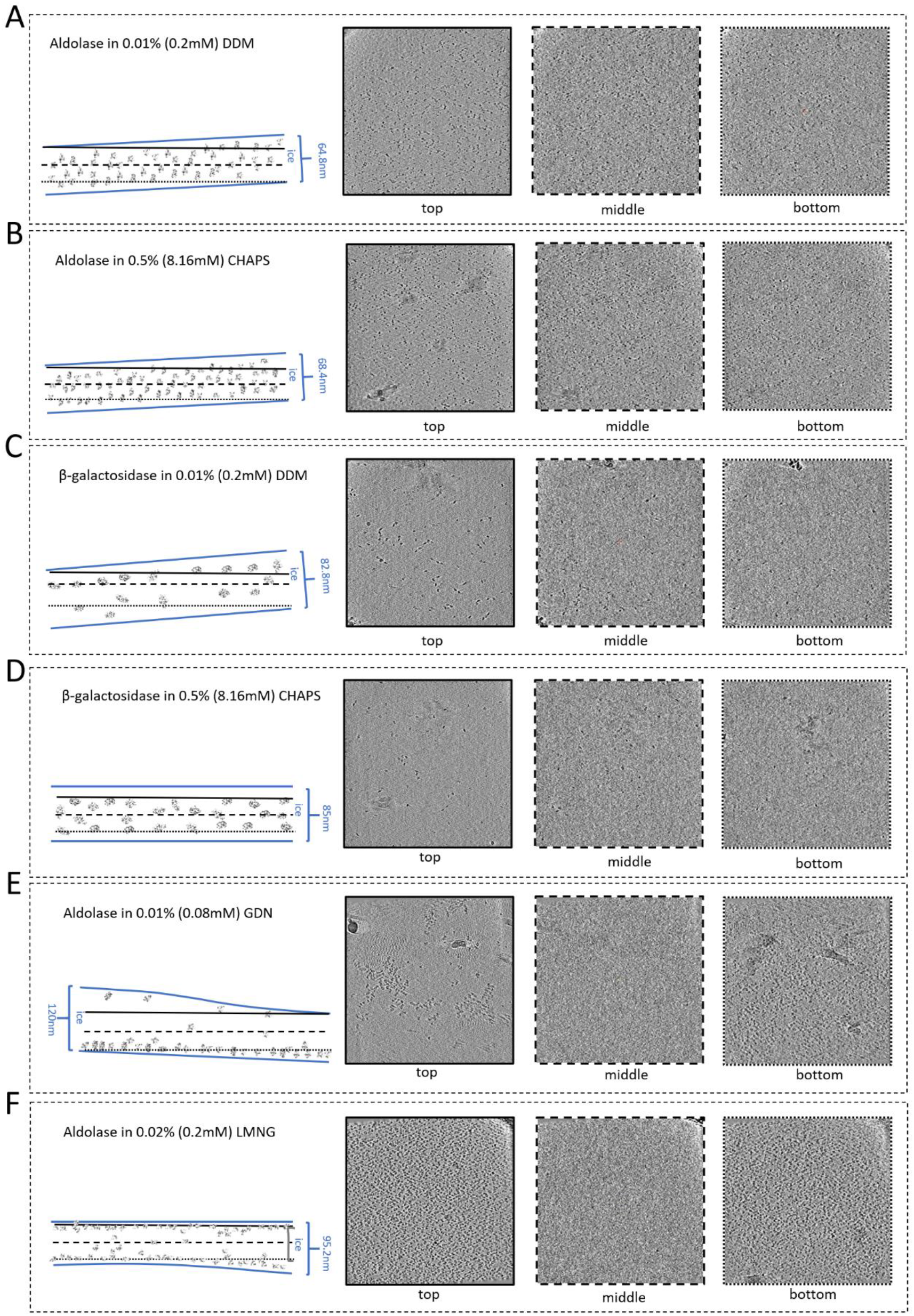
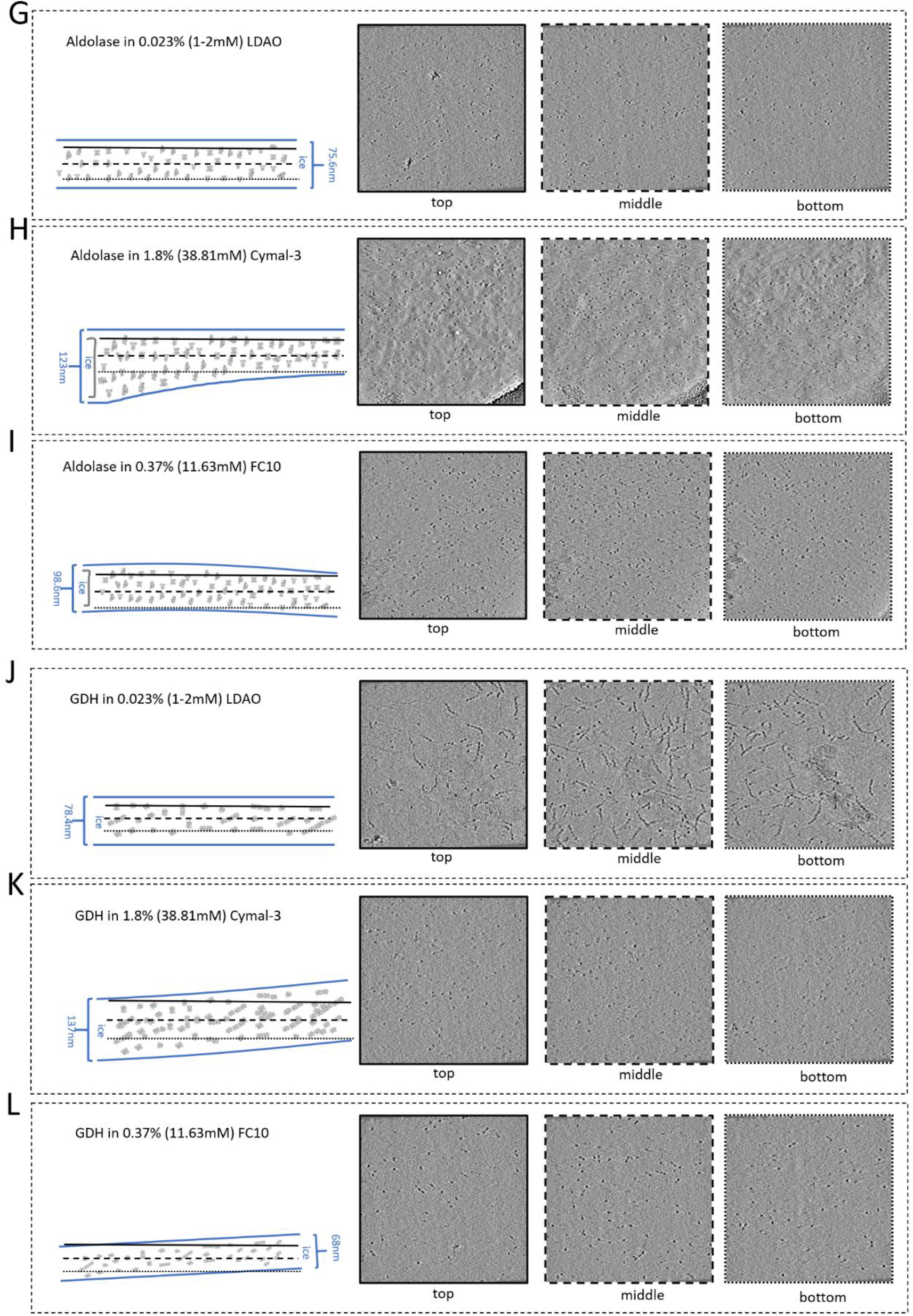
Slices of tomograms, showing variations in particle orientation of adsorbed and non-adsorbed particles of aldolase, β-galactosidase and GDH. (Left Panel) Schematic diagram of particle distribution in ice. Thickness of ice is indicated on the right. Different lines represent relative depths through tomographic reconstructions. (Right Panel) Top, middle and bottom tomographic cross-section.

**Figure S4.**
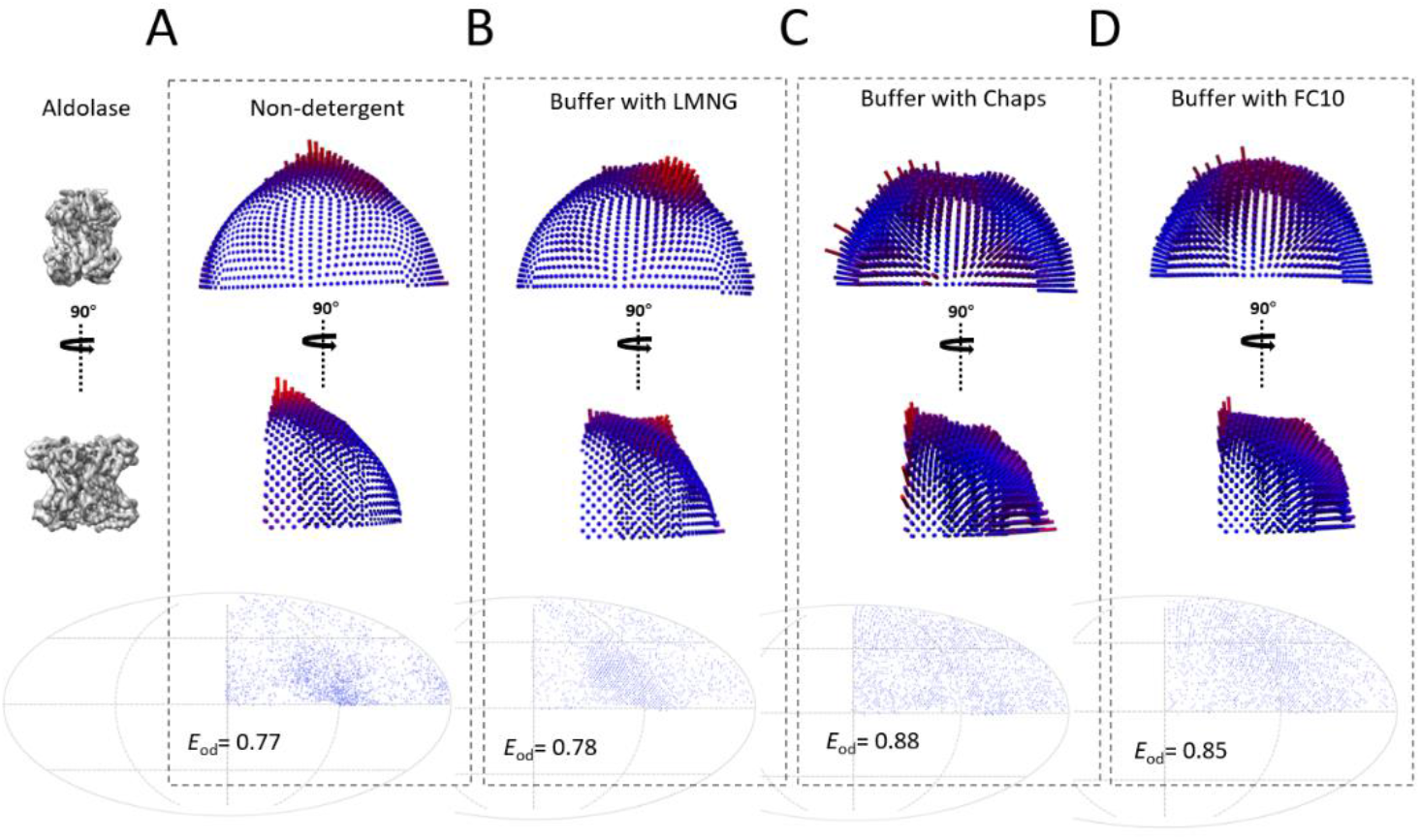
The aldolase orientation distribution in different detergents. (Top and Middle Panel) Orientation distribution sphere of the particles that contributed to the reconstruction presented corresponding to the views depicted in the left. The heights of the surface bars indicate the relative number of particles in a given orientation. (Bottom Panel) Orientation distributions plotted on a Mollweide projection. A: Angular orientation of aldolase in conventional buffer. B: Angular orientation of aldolase in buffer with 0.02% LMNG. C: Angular orientation of aldolase in buffer with 0.5% CHAPS. D: Angular orientation of aldolase in buffer with 0.37% FC10.

**Figure S5.**
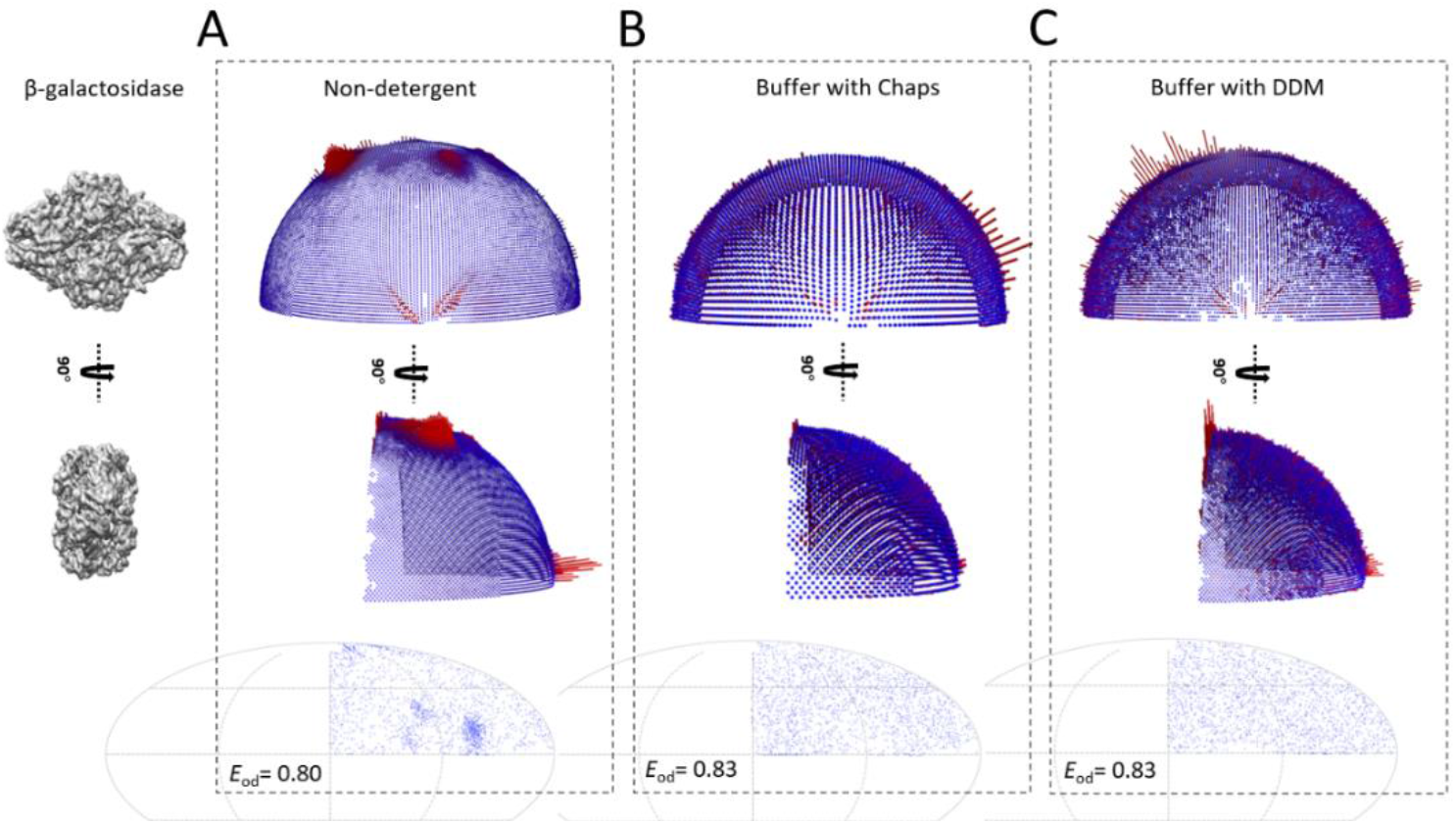
The β-galactosidase orientation distribution in different detergents. (Top and Middle Panel) Orientation distribution sphere of the particles that contributed to the reconstruction presented corresponding to the views depicted in the left. The heights of the surface bars indicate the relative number of particles in a given orientation. (Bottom Panel) Orientation distributions plotted on a Mollweide projection. A: Angular orientation of β-galactosidase in conventional buffer. B: Angular orientation of β-galactosidase in buffer with 0.5% CHAPS. C: Angular orientation of β-galactosidase in buffer with 0.01% DDM.

**Figure S6.**
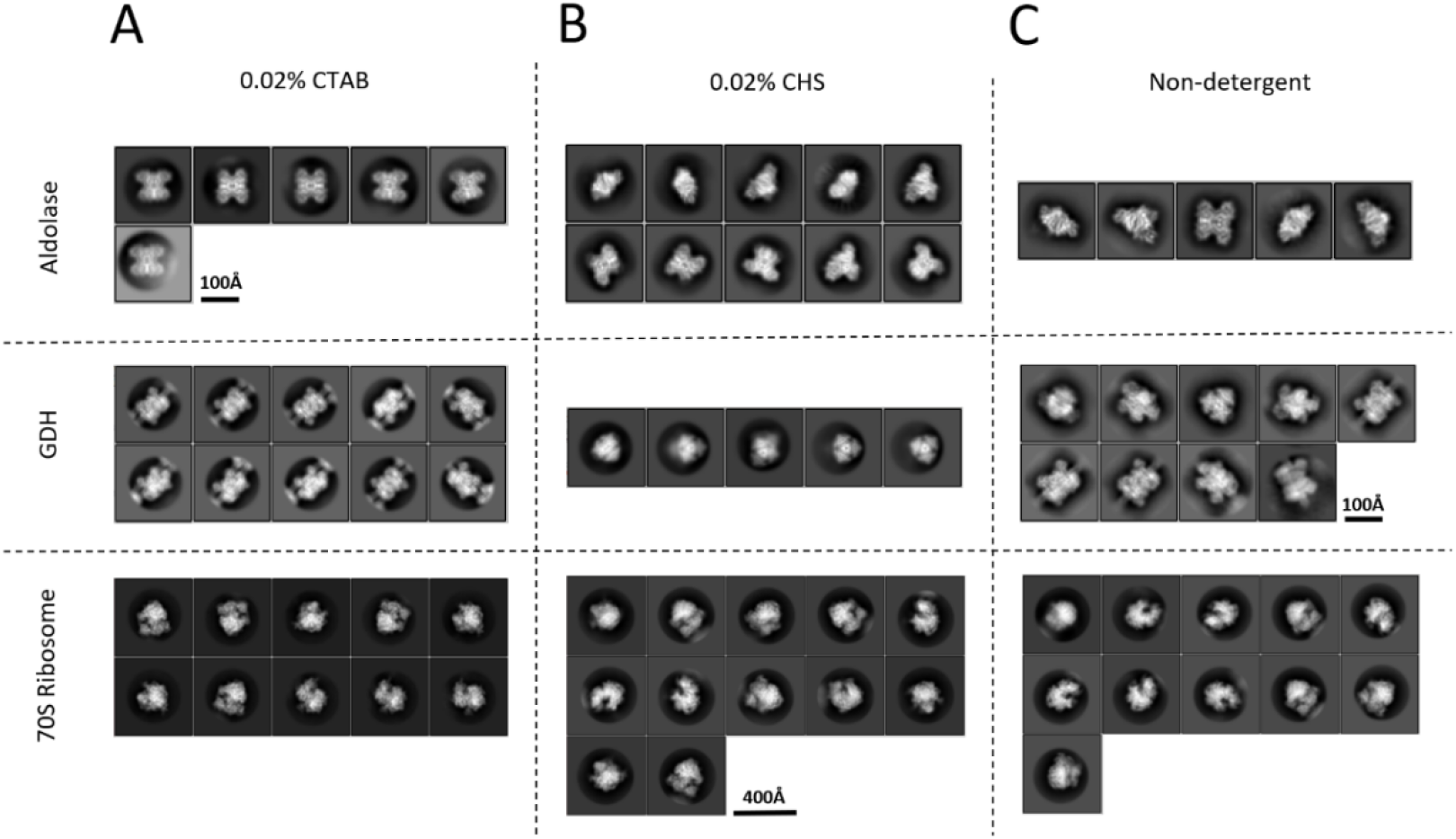
Comparison of single-particle cryo-EM 2D classifications obtained with different conditions. Representative the complete set of 2D class averages from aldolase, GDH and 70S Ribosomes grids prepared in buffer with 0.02%CTAB, buffer with 0.02%CHS or in standard buffer.

**Figure S7.**
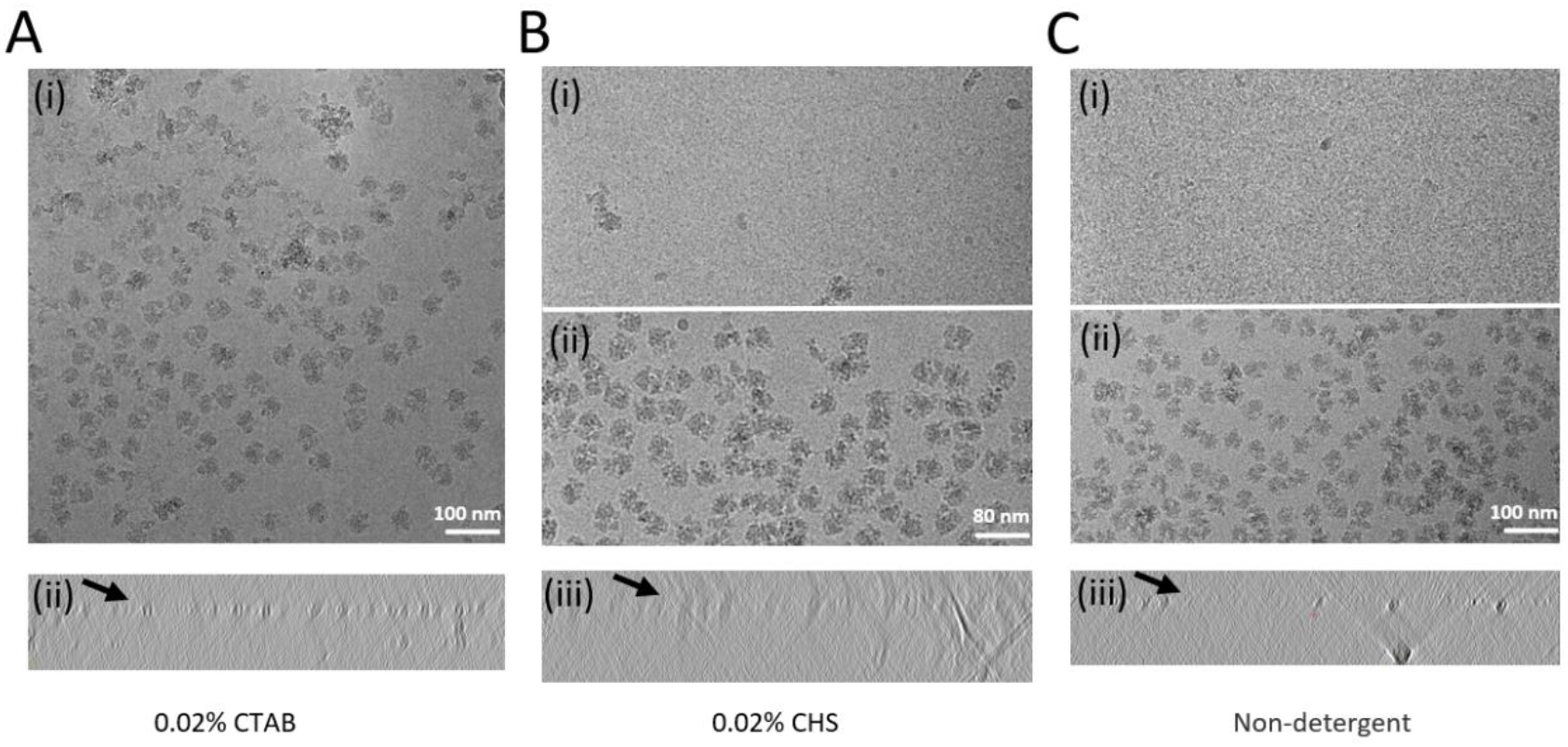
Representative high-magnification image and cross-sections from electron tomographic reconstructions of 70S Ribosomes. A: 80 nM Ribosomes in buffer with 0.02% CTAB prepared for cryo-EM. B: (i) 80 nM or (ii) 0.5 µM Ribosomes in buffer with 0.02% CHS prepared for cryo-EM. C: (i) 80 nM or (ii) 0.5 µM Ribosomes in standard buffer. (Bottom Panel) Cross-sections from electron tomographic reconstructions.

**Figure S8.**
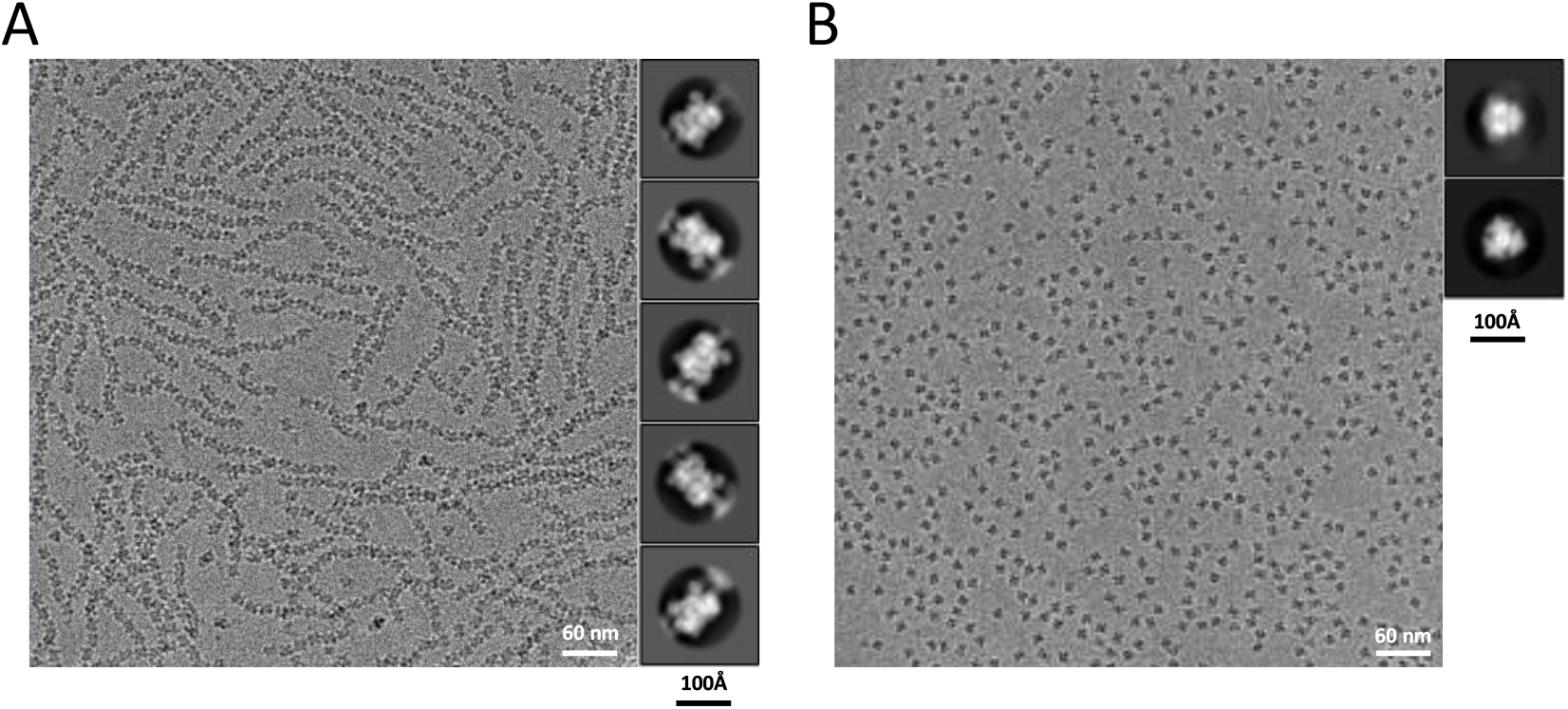
Representative high-magnification image and 2D class averages. A: Representative high-magnification image and 2D class averages of GDH in buffer with 0.02% DTAC. B: Representative high-magnification image and 2D class averages of GDH in buffer with 0.02% SLS.

**Figure S9.**
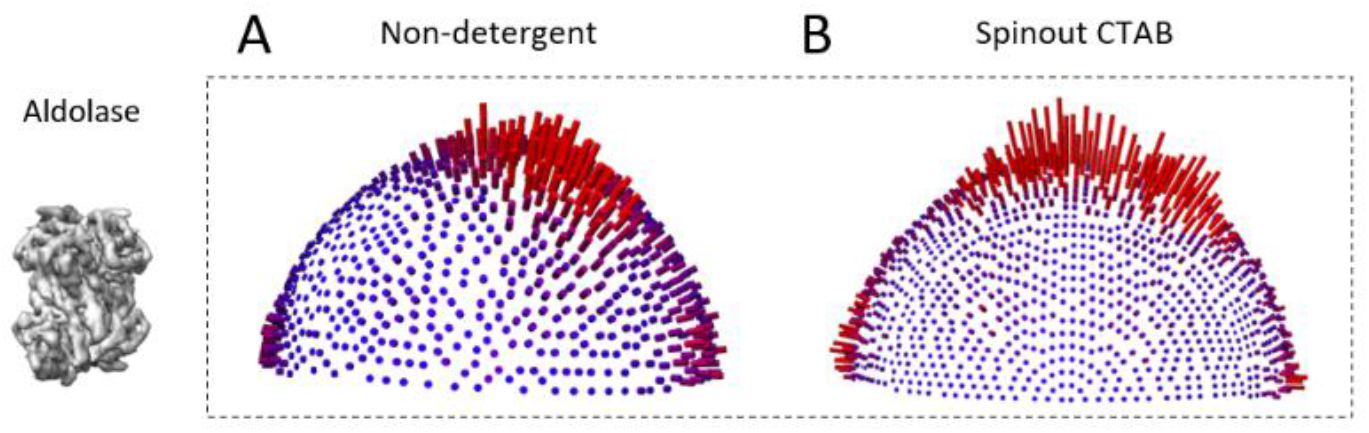
Orientation distribution in spinout CTAB sample resembles that in non-detergent sample.

## References

Brune D, Kim S J P O T N A O S 1993. Predicting protein diffusion coefficients. 90: 3835–3839.

Chaplin M J W 2009. Theory vs experiment: what is the surface charge of water. 1: 1–28.

Chen J, Noble A J, Kang J Y, et al. 2019. Eliminating effects of particle adsorption to the air/water interface in single-particle cryo-electron microscopy: Bacterial RNA polymerase and CHAPSO. 1: 100005.

Colloid E G J J O, Science I 2004. Surfactants and interfacial phenomena. 68: 347–347.

D’imprima E, Floris D, Joppe M, et al. 2019. Protein denaturation at the air-water interface and how to prevent it. 8: e42747.

Drzymala J, Sadowski Z, Holysz L, et al. 1999. Ice/water interface: Zeta potential, point of zero charge, and hydrophobicity. 220: 229–234.

Dubochet J, Adrian M, Chang J-J, et al. 1988. Cryo-electron microscopy of vitrified specimens. 21: 129–228.

Fan X, Wang J, Zhang X, et al. 2019. Single particle cryo-EM reconstruction of 52 kDa streptavidin at 3.2 Angstrom resolution. 10: 1–11.

Glaeser R M, Han B-G, Csencsits R, et al. 2016. Factors that influence the formation and stability of thin, cryo-EM specimens. 110: 749–755.

Glaeser R M, Han B-G J B R 2017. Opinion: hazards faced by macromolecules when confined to thin aqueous films. 3: 1–7.

Glaeser R M J C O I C, Science I 2018. Proteins, interfaces, and cryo-EM grids. 34: 1–8.

Grassucci R A, Taylor D J, Frank J J N P 2007. Preparation of macromolecular complexes for cryo-electron microscopy. 2: 3239.

Han Y, Fan X, Wang H, et al. 2020. High-yield monolayer graphene grids for near-atomic resolution cryoelectron microscopy. 117: 1009–1014.

Healthcare G, Healthcare F G 2007. Purifying challenging proteins principles and methods.

Klebl D P, Gravett M S, Kontziampasis D, et al. 2020. Need for speed: examining protein behavior during CryoEM grid preparation at different timescales. 28: 1238-1248. e1234.

Kontziampasis D, Klebl D P, Iadanza M G, et al. 2019. A cryo-EM grid preparation device for time-resolved structural studies. 6: 1024–1031.

Le maire M, Champeil P, Möller J V J B E B A-B 2000. Interaction of membrane proteins and lipids with solubilizing detergents. 1508: 86–111.

Naydenova K, Russo C J J N C 2017. Measuring the effects of particle orientation to improve the efficiency of electron cryomicroscopy. 8: 1–5.

Noble A J, Dandey V P, Wei H, et al. 2018a. Routine single particle CryoEM sample and grid characterization by tomography. 7: e34257.

Noble A J, Wei H, Dandey V P, et al. 2018b. Reducing effects of particle adsorption to the air–water interface in cryo-EM. 15: 793–795.

Plevka P, Battisti A J, Winkler D C, et al. 2012. Sample preparation induced artifacts in cryo-electron tomographs. 18: 1043.

Quinn P, Dawson R J B J 1970. An analysis of the interaction of protein with lipid monolayers at the air/water interface. 116: 671–680.

Ravelli R B, Nijpels F J, Henderikx R J, et al. 2019. Automated cryo-EM sample preparation by pin-printing and jet vitrification. 651208.

Rubinstein J L, Guo H, Ripstein Z A, et al. 2019. Shake-it-off: a simple ultrasonic cryo-EM specimen-preparation device. 75: 1063–1070.

Rubinstein J L J M 2007. Structural analysis of membrane protein complexes by single particle electron microscopy. 41: 409–416.

Sun F J C P B 2018. Orienting the future of bio-macromolecular electron microscopy. 27: 063601.

Takahashi M, Izawa E, Etou J, et al. 2002. Kinetic characteristic of bubble nucleation in superheated water using fluid inclusions. 71: 2174–2177.

Tan Y Z, Baldwin P R, Davis J H, et al. 2017. Addressing preferred specimen orientation in single-particle cryo-EM through tilting. 14: 793–796.

Taylor K A, Glaeser R M J J O S B 2008. Retrospective on the early development of cryoelectron microscopy of macromolecules and a prospective on opportunities for the future. 163: 214–223.

Vinothkumar K R, Henderson R J Q R O B 2016. Single particle electron cryomicroscopy: trends, issues and future perspective. 49.

Vos M R, Bomans P H, Frederik P M, et al. 2008. The development of a glove-box/Vitrobot combination: Air–water interface events visualized by cryo-TEM. 108: 1478–1483.

Young M, Carroad P, Bell R J B, et al. 1980. Estimation of diffusion coefficients of proteins. 22: 947–955.

